# Aligned basement membrane-modified collagen scaffolds for skeletal muscle tissue engineering

**DOI:** 10.64898/2026.07.11.736380

**Authors:** Ryann D. Boudreau, Geshani C. Bandara, Sarthak Pathak, Steven R. Caliari

## Abstract

Biomaterial scaffolds for repairing traumatic muscle injuries require restoration of both the anisotropic architecture and basement membrane extracellular matrix cues critical to normal muscle function. To address this need, we establish a collagen-glycosaminoglycan (CG) scaffold platform pairing an aligned pore microstructure, produced via directional freeze-drying, with basement membrane protein functionalization via carbodiimide crosslinking. Laminin and/or collagen IV are successfully tethered and retained within CG scaffolds over 7 days without significantly altering pore size or alignment, confirming stable protein functionalization and preservation of scaffold architecture. Human muscle progenitor cells show excellent viability and metabolic activity in all scaffold groups, with collagen IV functionalization significantly enhancing myotube number and fusion index. Toward establishing scaffold compatibility with non-myogenic support cells, we show that neural stem cells remain viable and metabolically active across all scaffold conditions. Overall, these findings highlight the combination of aligned scaffold architecture and collagen IV functionalization as potentially impactful for skeletal muscle tissue engineering.

## Introduction

The inherent regenerative capacity of skeletal muscle enables restoration of contractile tissue and function following minor injury^1^. However, traumatic injuries resulting in large scale loss of muscle fibers, extracellular matrix (ECM), and resident progenitor cells can overwhelm this regenerative capacity, resulting in volumetric muscle loss (VML)^2–6^. VML results in scarring rather than functional tissue regeneration, leaving patients with persistent functional deficits and long-term disability^7,8^. Current clinical management relies primarily on autologous muscle graft transfer supplemented with rehabilitative exercise^9–12^. Although the lost ECM infrastructure and composition are somewhat reinstated, this approach can lead to donor site morbidity^13,14^, mismatched dimensional and fiber type integration^15^, and incomplete functional recovery^16^.

Given the substantial burden of VML, biomaterial scaffold-based tissue engineering approaches have emerged as a promising alternative, offering a tunable platform to provide the structural and biological cues necessary to support muscle regeneration beyond the capabilities of current clinical options^17–19^. A critical design consideration for muscle-specific scaffolds is recapitulation of anisotropy, in which myofibers are aligned in parallel for directional force transmission and coordinated contraction^18,20^. Scaffolds with aligned pore structures have been shown to guide myotube orientation and enhance myogenic differentiation compared to isotropic controls^21,22^. Directional freeze-drying has been established as an effective fabrication method for producing scaffolds with controlled, anisotropic pore architecture, allowing for tunable pore size and alignment through modulation of freezing rate^23–27^. Type I collagen and glycosaminoglycans have been identified as particularly suitable base materials for these scaffolds^27–29^, and are already used in clinical settings to treat skin, tendon, and nerve injuries^30–33^. Prior work from our group demonstrated the utility of 3D, spatially aligned, acellular collagen-glycosaminoglycan (CG) scaffolds to induce *in vitro* myogenic differentiation in C2C12 myoblast cultures^27,34,35^ and partially support functional recovery in a rat VML model^36^. Although this platform sets the foundation for engineering anisotropic skeletal muscle, CG scaffolds lack the biochemical complexity of native skeletal muscle basement membrane proteins critical for supporting myogenic differentiation and functional recovery. This represents a key gap limiting their translational potential.

Myofiber basement membrane, composed primarily of laminin and type IV collagen (collagen IV), is a critical regulator of skeletal muscle regeneration, providing the adhesive and biochemical instructive cues governing muscle progenitor cell attachment, activation, and myogenic progression^37–40^. Laminin maintains the satellite cell (muscle stem cell) niche, supports progenitor cell self-renewal, and guides myogenesis^41–44^. Laminin 211 specifically facilitates cell survival, muscle integrity, and force transmission^45^. Collagen IV has been shown to promote differentiation and fusion of myoblasts by providing direct biochemical commitment cues to progenitor cells^46,47^. In VML injuries, the extensive loss of muscle tissue is accompanied by disruption and degradation of the basement membrane, severing these critical ECM-mediated signals and contributing to the fibrotic response observed clinically^17^. Reintroduction of laminin and collagen IV within a biomaterial scaffold therefore represents a rational strategy to restore the biochemical microenvironment necessary for productive myogenic cell adhesion and tissue regeneration.

Laminin incorporation into collagen-based scaffolds, through both chemical and physical crosslinking strategies, has been shown to support adhesion and proliferation across multiple cell types, including fibroblasts as well as nerve and endothelial cells^48–50^. Collagen IV functionalization has been explored in scaffold systems to improve cell-matrix interactions and tissue-specific function, with particular relevance to basement membrane reconstitution strategies^51,52^. In the context of skeletal muscle, collagen IV has been shown to promote myoblast differentiation and fusion in 2D cultures, establishing it as a functionally important pro-myogenic substrate^47^. However, the application of basement membrane protein-functionalized scaffolds specifically to skeletal muscle remains limited. Critically, prior functionalization studies have largely failed to characterize protein retention over extended culture periods, leaving open the question of whether observed cellular responses reflect sustained ECM signaling or an initial coating effect that degrades rapidly^53^. Furthermore, the impact of protein functionalization on the pore architecture and alignment of anisotropic scaffolds, which are properties essential for topographical guidance of myotube formation, has not been systematically evaluated. Taken together, these gaps highlight the need for a rigorous characterization of basement membrane protein functionalization in the context of an aligned scaffold system designed specifically for skeletal muscle tissue engineering.

Beyond driving myogenic cell bioactivity, skeletal muscle tissue engineering strategies must also support other key cell types, such as neural cells, critical for restoring innervation following injury^54^. Denervation of residual muscle fibers following VML further impairs contractile function independently of myofiber loss, and reinnervation is increasingly recognized as a critical and unaddressed component of functional VML recovery^55^. Neural stem cells (NSCs) represent a promising progenitor population for neural regeneration applications^56,57^ where aligned scaffold topography has been shown to direct organized neurite extension consistent with peripheral nerve organization^58–61^. Critically, laminin is among the most potent pro-neurogenic ECM components identified, supporting NSC adhesion, survival, and neuronal differentiation through integrin-mediated signaling^62–64^. However, the ability of aligned CG scaffolds (with or without basement membrane modification) to support NSC bioactivity has not been evaluated, representing a gap this work begins to address as a step toward establishing a scaffold platform supportive of the multiple cell types necessary for VML repair.

To address the identified limitations of current skeletal muscle tissue engineering scaffolds and ECM functionalization strategies, we present a strategy to fabricate aligned CG scaffolds functionalized with key basement membrane components of the skeletal muscle milieu (laminin, collagen IV, or their combination). Functionalized protein retention was characterized to confirm sustained biochemical presentation. Scaffold pore architecture and alignment were evaluated with and without basement membrane functionalization to confirm that the structural topographical cues essential for myotube guidance were preserved. Primary human muscle progenitor cells (hMPCs) were then seeded across all conditions to assess the influence of basement membrane protein composition on early myogenic state, metabolic activity, differentiation capacity, and myotube maturation and morphology. As a first step toward a neuromuscular co-culture platform, the same scaffold system was evaluated for its capacity to support primary neural stem cell adhesion and metabolic activity. Together, this work establishes a dual cell type validated platform for basement membrane-functionalized aligned scaffolds, advancing the translational foundation necessary for future neuromuscular tissue engineering and *in vivo* VML repair strategies.

## Methods

### Scaffold fabrication

Scaffolds were fabricated following a previously established protocol^27,35,65^. Briefly, 1.5 w/v% type I collagen from bovine Achilles tendon (Sigma-Aldrich) and 0.09 w/v% chondroitin sulfate sodium salt from bovine cartilage (Sigma-Aldrich) were dissolved in 0.05 M acetic acid. 63 mL of collagen solution was homogenized at 15,000 rpm using a recirculating chiller maintained at 4°C for 20 min, after which 7 mL of chondroitin sulfate solution was added, and homogenization continued for an additional 20 min. The suspension was degassed by vacuum and pipetted into cylindrical molds consisting of Teflon wells attached to a copper base. The suspension was directionally freeze-dried (Genesis 25L EL freeze dryer, SP Scientific) in molds at a freezing temperature of −10°C to produce scaffolds with anisotropic, aligned pore architecture. The resulting scaffold plugs measured 6.5 mm in diameter and 7.5 mm in height. Scaffolds were then placed in a vacuum oven at 105°C for 24 h to dehydrothermally crosslink.

### Scanning electron microscopy

Scanning electron microscopy (SEM) was performed on dehydrated CG scaffolds to visualize pore morphology and alignment. Scaffolds were sectioned using a scalpel blade to expose both the transverse and longitudinal planes, then mounted on aluminum stubs using carbon tape without sputter coating. Images were acquired using a Phenom XLG2 desktop SEM (Thermo Scientific) with a backscatter electron detector under low vacuum conditions (60 Pa) at an accelerating voltage of 10 kV.

### ECM protein functionalization

Scaffold plugs were sectioned transversely and longitudinally to produce semi-cylindrical scaffolds measuring 6.5 mm in diameter and 2.5 mm in height. Scaffolds were hydrated by immersion in 70% ethanol for 20 min on an orbital shaker at room temperature, then rinsed three times with phosphate buffered saline (PBS, ThermoFisher Scientific). Scaffolds were labeled with a 2 µM solution of AlexaFluor 568 NHS ester in PBS (1 mL/scaffold, ThermoFisher Scientific) for 20 min on an orbital shaker at room temperature. Scaffolds were then washed twice with PBS prior to coating and crosslinking.

Scaffolds were incubated in 235 µL of their respective protein solutions for 2 h at 37°C on an orbital shaker at 100 rpm: laminin alone (10 μg/cm^2^, human laminin 211, Biolamina), collagen IV alone (10 μg/cm^2^, human collagen IV, Fisher Scientific), or a combination of both proteins (10 μg/cm^2^ each). Protein concentrations were calculated based on the surface area of the porous scaffold^66^. Uncoated CG scaffolds were incubated with PBS under identical conditions. All scaffolds were subsequently crosslinked using 1-ethyl-3-(3-dimethylaminopropyl) carbodiimide hydrochloride (EDC, Sigma Aldrich) and N-hydroxysulfosuccinimide (NHS, Sigma Aldrich) at a 5:2:1 molar ratio^67^, relative to the carboxylic acid content of the collagen scaffold, for 50 min at 37°C on an orbital shaker at 100 rpm. Scaffolds were then sterilized in 70% ethanol for 20 min and rinsed three times with sterile PBS.

### ECM protein retention analysis

ECM-coated scaffolds were incubated in cell growth media at 37°C to assess protein retention over 7 days, with scaffolds collected at day 0 and day 7 (*N* = 3 per group per timepoint). Cell growth media consisted of Dulbecco’s Modified Eagle Medium/Nutrient Mixture F-12 (DMEM/F12, 4.5g/L glucose, ThermoFisher Scientific) supplemented with 18% fetal bovine serum (FBS, Gibco), 50 µg/mL gentamicin (ThermoFisher Scientific), 10 ng/mL human epidermal growth factor (hEGF, ThermoFisher Scientific), 1 ng/mL human fibroblast growth factor (hFGF, ThermoFisher Scientific), 10 µg/mL Insulin-Transferrin-Selenium (ThermoFisher Scientific), and 0.4 µg/mL dexamethasone (ThermoFisher Scientific). Scaffolds were fixed in 10% formalin for 20 min at room temperature and then rinsed twice with PBS. Scaffolds were permeabilized in 0.1% Triton X-100 (Sigma Aldrich) for 20 min at room temperature on an orbital shaker, then blocked in 3% bovine serum albumin (BSA, Sigma Aldrich) overnight at 4°C. ECM proteins were detected using the following primary antibodies: anti-laminin (anti-laminin 1+2, rabbit polyclonal, Abcam, 1:200) and anti-collagen IV (anti-collagen IV antibody, mouse monoclonal, Millipore Sigma, 1:200), incubated overnight at 4°C on an orbital shaker. Secondary antibodies included AlexaFluor 488-conjugated goat anti-rabbit and AlexaFluor 647-conjugated goat anti-mouse, each at 1:400 dilution, incubated for 2 h at room temperature on an orbital shaker. Scaffolds were rinsed twice with PBS following each antibody incubation. Samples were imaged using a Cytation C10 confocal microscope with a z-stack depth of 50 µm and step size of 10 µm. Image acquisition parameters were held constant across all samples and time points to enable quantitative fluorescence intensity comparisons. Maximum intensity z-projections were generated and imported into CellProfiler for quantification of mean fluorescence intensity per scaffold, with values compared between day 0 and day 7 to assess protein retention. FIJI (ImageJ) was used to merge fluorescence channels for qualitative visualization of protein distribution within the scaffold.

### Pore morphology assessment

Pore architecture of the ECM-coated scaffolds was evaluated using confocal images of the transverse and longitudinal planes, acquired using the immunofluorescence staining protocol described above. Quantification of pore diameter, area, aspect ratio, and orientation was performed using a custom Python pipeline applied to individual z-stack TIFF files. Briefly, fluorescence channels were merged to produce a composite image of the scaffold architecture. The sharpest focal plane within each z-stack was identified using a Laplacian-based focus metric, in which the variance of the Laplacian was computed for each slice and the slice with the highest variance was selected for analysis. Pore segmentation masks were manually drawn over individual pores using a Python-based interactive viewer built with Matplotlib and NumPy. A minimum of 250 pores were segmented per scaffold (*N* = 3 scaffolds per group). The skimage regionprops package was used to extract the major axis, minor axis, and area from each segmentation mask. Structure tensor analysis was used to obtain the orientation from each image. Aspect ratios were computed by dividing the major axes by the minor axes. Pore diameter was calculated as the root mean square of the major and minor axes of the best-fit ellipse, providing a diameter estimate that accounts for the pore ellipticity:

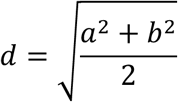

where *a* is the major axis length and *b* is the minor axis length. Segmentation artifacts smaller than 100 pixels (equivalent to 34 µm at the imaging magnification used) were excluded from analysis. The circular axial mean orientation was calculated and used as a reference angle for normalization of individual pore orientations. Normalized orientations were binned in 5° increments and plotted as rose diagrams to visualize pore alignment per group.

### Cell culture

Primary hMPCs were obtained from Dr. George Christ’s lab at the University of Virginia from a 21-year-old female donor. Primary adult rat hippocampal neural stem cells (NSCs, SCR022, Millipore Sigma), isolated from adult female Fisher 344 rats aged 43-55 days, were obtained commercially. Cryopreserved cells were thawed rapidly in a 37°C water bath, diluted dropwise into pre-warmed growth media, centrifuged at 200 × g for 5 min, and resuspended in fresh growth media prior to initial plating. HMPCs were used between passages 2 and 3 and NSCs were used at passage 2. NSC growth media consisted of DMEM/F12 (without L-glutamine, 4.5g/L glucose, ThermoFisher Scientific) supplemented with 2% Stempro neural supplement (ThermoFisher), 18% FBS (Gibco), 50 µg/mL gentamicin (ThermoFisher Scientific), 10 ng/mL human epidermal growth factor (hEGF, ThermoFisher Scientific), 1 ng/mL human fibroblast growth factor (hFGF, ThermoFisher Scientific), 10 µg/mL insulin-transferrin-selenium (ThermoFisher Scientific), and 0.4 µg/mL dexamethasone (ThermoFisher Scientific). Both cell types were cultured in T75 flasks coated with Matrigel (0.083 mg/mL, Corning), in their respective growth media described above, and maintained at 37°C in a humidified 5% CO_2_ incubator. HMPCs underwent media changes every 3 days while NSCs underwent 50% media volume changes every 3 days. Upon reaching 80% confluency, media was aspirated, and cells were rinsed with PBS. Cells were detached by incubation in Accutase solution (Sigma-Aldrich) for 3 min at 37°C, with gentle agitation of the flask to facilitate detachment. Detachment was quenched by addition of double the volume of cell growth media, and the cell suspension was centrifuged at 200 × g for 5 min. The resulting pellet was resuspended in growth media and cells were counted using a Countess automated counter (ThermoFisher) prior to seeding.

### Scaffold cell seeding

Prior to cell seeding, scaffolds were incubated in their respective cell growth media at 37°C on an orbital shaker at 100 rpm for 30 min to ensure complete hydration. Scaffolds were seeded with 1 mL of cell suspension at a concentration of 200,000 cells/mL (200,000 cells per scaffold, *N* = 3-6 per group) in an ultra-low attachment 24-well plate and were incubated overnight at 37°C on an orbital shaker at 100 rpm to promote uniform cell distribution throughout the scaffold volume. Scaffolds were then transferred to 1 mL of fresh growth media, replaced every 2 days. For muscle differentiation experiments, growth media was replaced with differentiation media after 2-3 days of culture. Muscle differentiation media consisted of DMEM/F12 (4.5 g/L glucose) supplemented with 10% FBS, 50 µg/mL gentamicin, and 10 µg/mL insulin-transferrin-selenium, replaced every 2 days. Muscle differentiation cultures were maintained for a total of 3 days before fixation and analysis. For neural differentiation experiments, growth media was replaced with differentiation media after 2 days of culture. Neural differentiation media consisted of Neurobasal medium (ThermoFisher) supplemented with 2% B-27 Supplement (ThermoFisher), 1% L-glutamine (GlutaMAX-1, Gibco), 10% FBS, 50 µg/mL gentamicin, and 10 µg/mL insulin-transferrin-selenium, with 50% media volume changes every 2 days. Neural differentiation cultures were maintained for a total of 6 days before fixation and analysis.

### Cell viability assessment

Cell viability was qualitatively assessed on day 6 of culture using a Live/Dead cytotoxicity assay (ThermoFisher Scientific). A staining solution was prepared by combining 5 µL calcein AM, 20 µL ethidium homodimer-1, and 10 mL PBS. Media was aspirated and scaffolds were rinsed with PBS before the addition of 1 mL staining solution per well. Scaffolds were incubated for 20 min at room temperature on an orbital shaker, protected from light. Scaffolds were rinsed with PBS and imaged using the Cytation C10 confocal microscope at 20x magnification. Maximum intensity z-projections were generated from z-stacks of 100 µm depth acquired at 10 µm step intervals. Scaffolds used for Live/Dead imaging were not previously stained with AlexaFluor 568 NHS ester.

### Metabolic activity assay

Metabolic activity of cells seeded on CG scaffolds was non-destructively assessed using alamarBlue (ThermoFisher Scientific). The same scaffolds were measured at each time point. Each scaffold was incubated in 1 mL of 10% alamarBlue reagent diluted in cell growth media at 37°C on an orbital shaker for 70 min, protected from the light. Three 100 μL aliquots of conditioned media per scaffold were transferred into a 96-well plate as technical replicates. Fluorescence was measured using a Tecan Infinite M Plex multimode plate reader at excitation and emission values of 550 nm and 590 nm respectively. AlamarBlue reagent blanks were prepared and measured identically at each timepoint, and blank fluorescence values were subtracted from corresponding experimental measurements. Blank-subtracted fluorescence values were plotted as a function of culture day for each scaffold group to assess relative metabolic activity over time (*N* = 3 scaffolds per group).

### Immunofluorescence staining

Scaffolds were fixed in 10% neutral buffered formalin for 20 min at room temperature and rinsed twice with PBS. Scaffolds were permeabilized and blocked as described above. For assessment of myotube formation, muscle cell-laden scaffolds were incubated overnight at 4°C on an orbital shaker with anti-myosin heavy chain primary antibody (MHC, mouse monoclonal, eBiosciences, 1:400). For assessment of neuronal differentiation, neural cell-laden scaffolds were incubated overnight at 4°C on an orbital shaker in anti-β-III tubulin (rabbit polyclonal, Abcam, 1:400) primary antibody. Following primary antibody incubation, all scaffolds were rinsed twice with PBS. Muscle cell-laden scaffolds were incubated for 2 h at room temperature on an orbital shaker with AlexaFluor 488-conjugated goat anti-mouse secondary antibody (ThermoFisher, 1:800) and Alexa Fluor 647-conjugated phalloidin (ThermoFisher, 1:800), for actin cytoskeleton visualization. Neural cell-laden scaffolds were incubated under identical conditions with AlexaFluor 647-conjugated donkey anti-rabbit secondary antibody (1:800) secondary antibody. All scaffolds were rinsed twice with PBS before nuclear labeling with 4’,6-diamidino-2-phenylindole (DAPI, ThermoFisher, 1:1000) for 5 min at room temperature, followed by two final PBS rinses. Scaffolds were imaged on transverse and longitudinal planes using a Cytation C10 confocal microscope at 20x magnification. Z-stacks of 100 µm depth were acquired at 10 µm step intervals and maximum intensity z-projections were generated for analysis (muscle: *N* = 6 scaffolds per group; neural: *N* = 3 scaffolds per group).

### Myotube formation and morphometric analysis

Myotube formation and morphology were assessed following 3 days of differentiation media culture. Maximum intensity z-projections of DAPI and MHC fluorescence channels were processed using Omnipose to segment both nuclei and MHC+ myotube regions. Segmentation masks were manually reviewed and corrected in Python to remove artifacts and ensure accuracy. The skimage regionprops package was used to extract major axis length, minor axis length, and area from each mask. Structure tensor analysis was used to obtain the pixel orientations from each image. Myotube masks were filtered by size based on inspection of area distributions, with objects below 688 µm^2^ (1000 pixels) excluded as segmentation artifacts. The number of nuclei masks overlapping each MHC+ myotube mask was quantified to determine nuclei per myotube. Fusion index was calculated as the number of nuclei masks overlapping an MHC+ mask divided by the total number of nuclei masks. Three fields of view were analyzed per scaffold (*N* = 6 scaffolds per group).

### Statistical analysis

Statistical analysis was performed doing one-way or two-way ANOVA with pairwise comparisons between groups performed using estimated marginal means with Tukey adjustment for multiple comparisons using the emmeans package in R statistical software. Significance was defined as p values < 0.05. Experiments were conducted with *N* = 3-6 scaffolds per group as biological replicates and *n* = 3-4 fields of view per scaffold as technical replicates. Data are presented as mean ± standard deviation with individual data points shown.

## Results and Discussion

### Directional freeze-drying produces aligned porous CG scaffolds suitable for ECM functionalization

The CG scaffold system utilized for this study was fabricated via directional freeze-drying of a suspension consisting of type I collagen and chondroitin sulfate in a Teflon/copper mold (**Fig. 1a**). The thermal gradient established by the copper base drives directional ice crystal growth and produces anisotropic pore architecture upon sublimation (**Fig. 1b**). Sectioning of the dehydrated scaffold plugs along either longitudinal or transverse planes revealed distinct pore morphologies (**Fig. 1c**). SEM imaging of the transverse plane, oriented perpendicular to the freezing direction, revealed a randomized circular arrangement of pores without a preferential orientation (**Fig. 1d**). In contrast, the longitudinal plane, oriented parallel to the freezing direction, showed elongated channels aligned along the axis of the ice crystal propagation (**Fig. 1e**). This anisotropic architecture recapitulates the parallel organization of the native myofiber bundles and provides the topographical framework for directional guidance of cell outgrowth.

**Figure 1.**
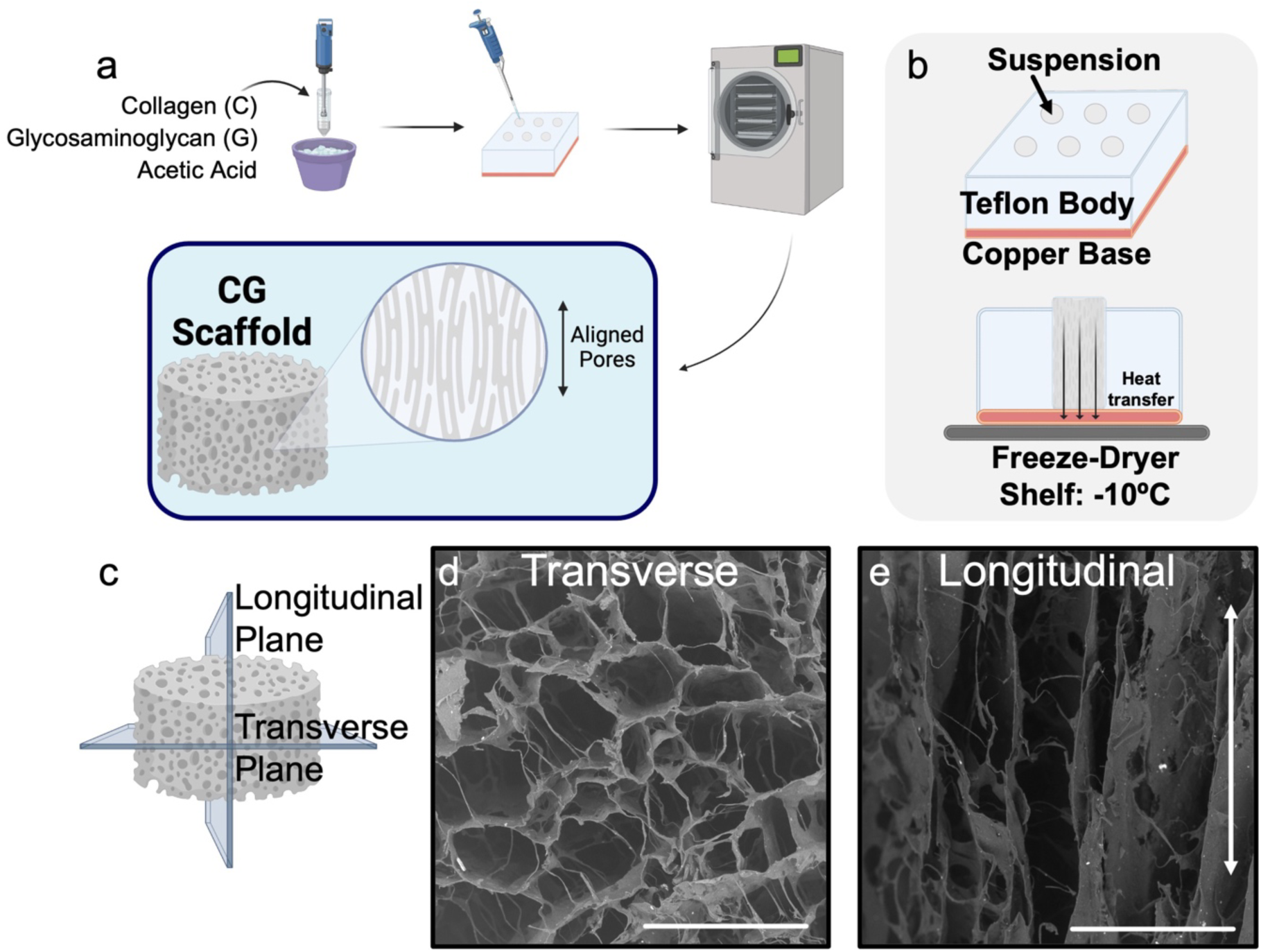
Directionally freeze-dried collagen-glycosaminoglycan (CG) scaffolds recapitulate muscle anisotropy. (a) Scaffold fabrication involves homogenization of type I collagen and chondroitin sulfate in acetic acid to form a suspension, followed by freeze-drying to generate porous scaffold plugs. (b) Directional freezing is achieved by lyophilizing the suspension in a Teflon mold with a copper base, creating a thermal gradient that induces anisotropic pore formation. (c) Resulting scaffolds can be analyzed in both the longitudinal (parallel to the freezing direction) and transverse planes. Scanning electron microscope (SEM) images reveal a randomized pore structure in the (d) transverse plane, and a highly aligned, anisotropic structure in the (e) longitudinal plane, highlighting differences in pore structure. Scale bars: 300 µm.

### Laminin and collagen IV remained functionalized to CG scaffolds over 7 days

To enhance the biological relevance of CG scaffolds for skeletal muscle tissue engineering, subsequent experiments investigated scaffold functionalization with basement membrane ECM proteins laminin 211 and/or collagen IV. CG scaffolds were coated with laminin and/or collagen IV via EDC/NHS covalent tethering. Functionalization was assayed via antibody-based confocal fluorescence microscopy of modified scaffolds. Laminin and collagen IV fluorescence was detected throughout the scaffolds in all respective coating groups at day 0, confirming successful ECM protein functionalization prior to cell culture (**Fig. 2a-d**). Fluorescence intensity was maintained through 7 days of incubation across all groups (**Fig. 2e-h**), demonstrating that EDC/NHS crosslinking achieved stable protein tethering over the experimental period. Quantitative analysis of maximum fluorescence intensity confirmed no significant decrease in laminin signal between day 0 and day 7 for the laminin alone or combined coating groups (**Fig. 2i**). There was also no significant decrease in collagen IV signal for the collagen IV alone or combined groups (**Fig. 2j**). Notably, the combined laminin/collagen IV (Lam/Col) group exhibited significantly lower collagen IV fluorescence intensity compared to the collagen IV alone group at both timepoints. This could suggest that simultaneous application of both proteins may result in laminin limiting collagen IV binding due to steric hindrance and/or non-substrate binding^68–70^. Scaffold backbone fluorescence intensity remained stable over time, confirming that observed changes in ECM protein signal reflect true protein retention rather than scaffold autofluorescence artifacts (**Fig. S1**). Z-stack analysis further revealed that both laminin and collagen IV fluorescence was detected not just at the scaffold surface, but throughout the full 50 µm image depth at both time points (**Fig. S1**).

**Figure 2.**
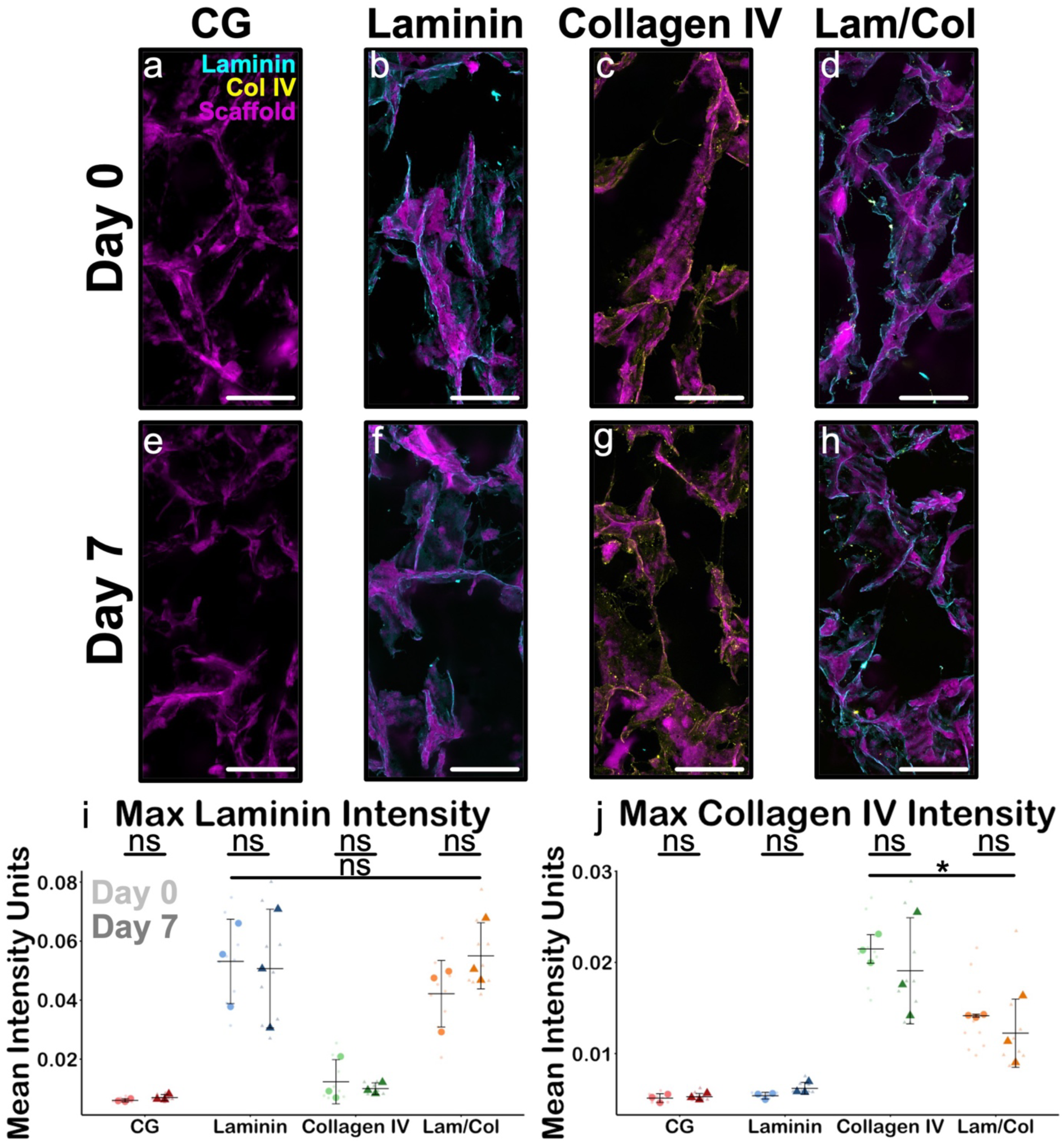
ECM proteins remain stably associated within scaffolds over the experimental period. Representative confocal images of scaffolds at (a-d) day 0 and (e-h) day 7 show laminin (cyan), collagen IV (yellow), and scaffold backbone (magenta) present within their respective experimental groups. (i) Quantification of fluorescence intensity demonstrates no significant reduction in laminin signal across groups between day 0 and day 7, or in (j) collagen IV signal in the collagen IV group, indicating stable retention of ECM proteins within the scaffolds. However, there is a significant decrease in collagen IV signal in the laminin/collagen IV group at both time points compared to the collagen IV alone group. Scale bars: 200 µm. *N* = 3 scaffolds per group. *: p < 0.05.

### Pore morphology and organization remained consistent across all ECM-modified groups

Having confirmed the ability to stably coat CG scaffolds with basement membrane ECM proteins, subsequent experiments evaluated whether functionalization altered CG scaffold pore architecture. Confocal imaging of fluorescently-labeled scaffolds revealed distinct pore morphologies in the longitudinal versus transverse planes for all experimental groups. The longitudinal plane exhibited elongated, parallel pore channels oriented along the freeze-drying axis (**Fig. 3a-d**), while the transverse plane showed a non-preferential, roughly isotropic pore arrangement (**Fig. 3e-h**). Quantitative analysis confirmed that pore diameter did not significantly differ between coating groups in either plane (**Fig. 3i**). Mean longitudinal pore diameters ranged from 160-219 µm and mean transverse pore diameters spanned 146-179 µm across groups, values consistent with the reported optimal pore size range of 100-300 µm for skeletal muscle cell infiltration^27,66^. Aspect ratios followed a similar pattern, with longitudinal values ranging from 2.07 to 2.50 and transverse values from 1.89 to 2.23 (**Fig. 3j**), with longitudinal aspect ratios trending slightly higher than transverse, consistent with the elongated pore morphology observed in longitudinal plane images as a result of directional freezing. Pore orientation analysis revealed that longitudinal plane pores showed a higher frequency of alignment at the reference orientation compared to transverse plane pores across groups, with longitudinal peak frequencies of 7.3-9.8% compared to 4.9-7.4% in the transverse plane (**Fig. 3k**). Taken together, these findings demonstrate that ECM protein functionalization does not substantially alter the pore size, aspect ratio, or alignment of the CG scaffolds, preserving the topographical architecture critical for directional guidance of muscle progenitor cells.

**Figure 3.**
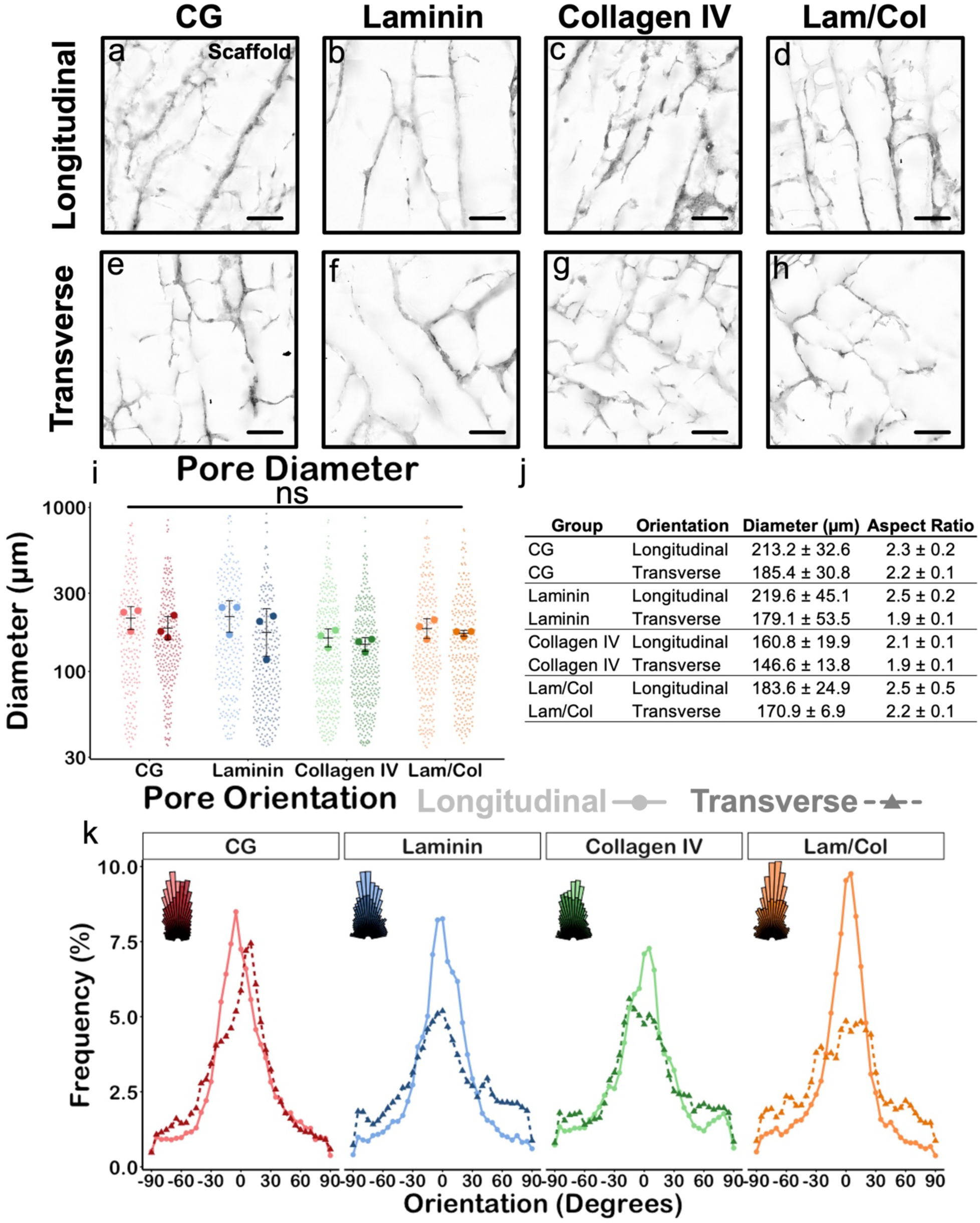
ECM protein functionalization does not alter scaffold pore morphology. Representative confocal images of the scaffold (a-d) longitudinal plane and (e-h) transverse plane mimicking anisotropic skeletal muscle architecture. (i) Quantitative analysis indicates no significant differences in pore diameter across experimental groups, showing that ECM functionalization preserves pore size and morphology. (j) Summary of pore diameter and aspect ratio values for each group show larger values for the longitudinal plane. (k) Analysis of pore orientation reveals anisotropic alignment in the longitudinal planes of each group. Scale bars: 200 µm. *N* = 3 scaffolds. *: p < 0.05.

### hMPCs are viable and metabolically active in all ECM-modified scaffold groups

Having confirmed maintenance of pore size and alignment in ECM-functionalized CG scaffolds, subsequent experiments assessed cellular response. *In vitro* validation of biomaterial scaffolds for skeletal muscle regeneration has relied predominantly on murine myoblast cell lines, most commonly C2C12s, due to their ease of culture, high proliferative capacity, and reliable differentiation behavior^19,22,71,72^. However, these immortalized cell lines differ substantially from primary hMPCs in their dependence on microenvironmental cues, their myogenic differentiation potential, and their sensitivity to ECM composition^72–75^. Species-specific differences between murine and human myogenic cells further limit the direct translation of findings from C2C12-based studies to human clinical applications^76,77^. Primary hMPCs recapitulate the endogenous cells that would respond to an implanted scaffold in a clinical setting, providing a more rigorous and translationally meaningful platform for scaffold validation. While their use introduces greater biological variability and more demanding culture requirements, these characteristics reflect the complexity of the human muscle regenerative environment and strengthen the clinical relevance of the experimental findings.

Scaffolds seeded with hMPCs were cultured for 3 days in growth media and 3 days in myogenic differentiation media. All scaffold groups supported increasing hMPC metabolic activity over the culture period, with fluorescence values rising significantly from day 1 through day 4 before plateauing between days 4 and 6 (**Fig. 4a**). This plateau in metabolic activity is consistent with hMPCs exiting the proliferative cycle and committing to terminal myogenic differentiation, as post-mitotic myotubes exhibit reduced metabolic turnover compared to actively dividing myoblasts. Importantly, no significant differences in metabolic activity were detected between groups at any time point, indicating that laminin, collagen IV, and their combination did not adversely affect hMPC metabolic health relative to uncoated CG scaffolds. These findings are consistent with prior work demonstrating that EDC/NHS-crosslinked ECM coatings on collagen-based scaffolds maintain cytocompatibility^78^. Live/dead imaging on day 6 qualitatively confirmed high cell viability across all scaffold groups (**Fig. 4b-e**). Supplemental analysis of myogenic state at 24 hours post-seeding revealed comparable proportions of Pax7+ and MyoD+ cells across all experimental groups (**Fig. S2**), suggesting that ECM functionalization did not substantially shift cell adhesion or early myogenic commitment at this time point. Collectively these findings confirm that all ECM-functionalized scaffold groups supported hMPC metabolic activity and viability equivalently, establishing a healthy cellular baseline for subsequent differentiation analysis.

**Figure 4.**
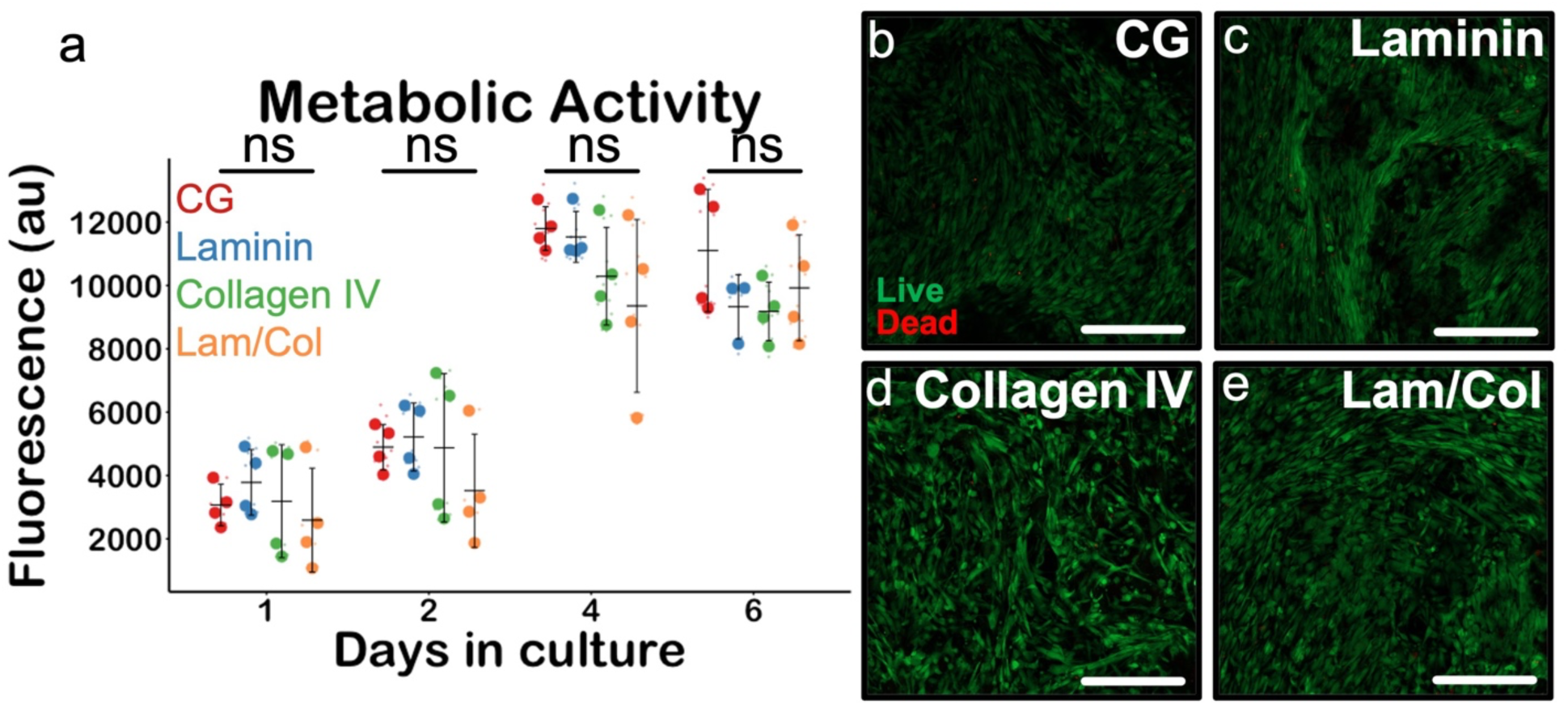
All ECM functionalization conditions support myoblast metabolic health. (a) Fluorescence measurements taken from alamarBlue assay show consistent cell metabolic activity between groups. *N* = 3 scaffolds. (b-e) Representative Live/Dead images after 6 days of culture suggest positive cell viability and homogenous distribution of cells in each group. Scale bars: 200 µm.

### Collagen IV-functionalized CG scaffolds support increased myogenic differentiation

Fluorescence imaging of all scaffold groups at the differentiation endpoint revealed robust cell coverage as indicated by DAPI and F-actin staining, with clearly defined MHC+ myotubes present across all conditions (**Fig. 5a-d**). Examination of isolated MHC channels confirmed multinucleated myotube formation with morphological evidence of alignment along a similar axis in all groups (**Fig. 5e-h**). Quantification of total myotube area and myotube count revealed comparable scaffold coverage across most groups, with a significant reduction in the laminin/collagen IV group compared to the collagen IV group (**Fig. 5i-j**). This suggests that overall cell engraftment and myotube formation capacity were maintained across single protein coating conditions but were diminished in the combined coating group.

**Figure 5.**
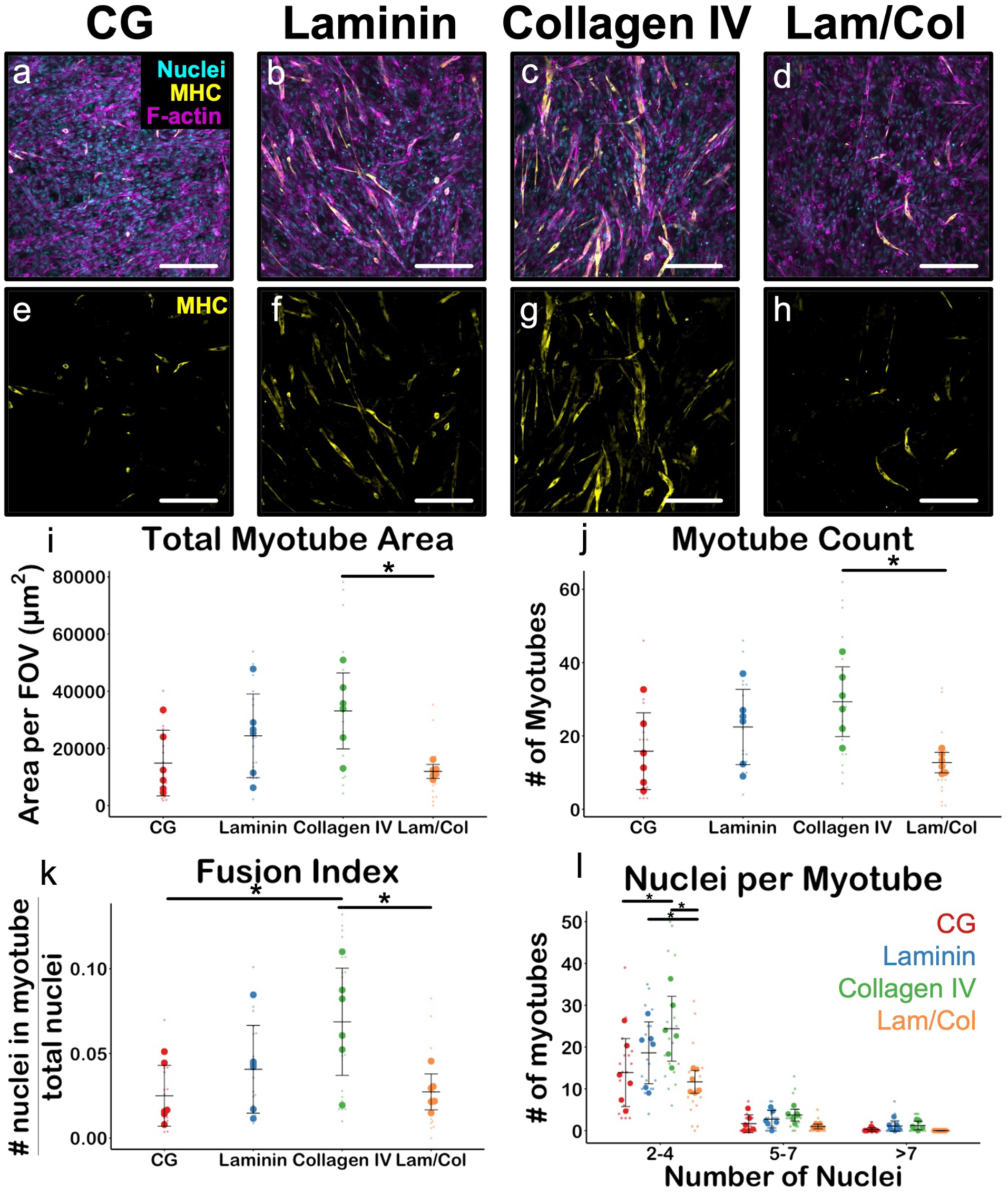
Collagen IV-functionalized scaffolds support increased myogenic differentiation. (a-d) Representative fluorescence images of hMPCs cultured on ECM-functionalized scaffolds stained for myosin heavy chain (MHC, yellow), F-actin (magenta), and nuclei (cyan). (e-h) MHC-only channel images demonstrate myotube formation. (i) Quantification of total myotube area per field of view (FOV) and (j) average number of myotubes per group shows significantly increased levels in the collagen IV group compared to the laminin/collagen IV group. (k) The collagen IV group also showed a significantly increased fusion index compared to the CG and laminin/collagen IV groups. (l) Graph depicting nuclei per myotube indicates a trend toward increased maturation in the collagen IV group relative to the other conditions. Scale bars: 200 µm. *N* = 6 scaffolds. *: p < 0.05.

Fusion index analysis revealed that the collagen IV group achieved significantly greater myoblast fusion compared to both the laminin/collagen IV and uncoated CG groups (**Fig. 5k**), indicating that collagen IV functionalization actively promotes myogenic fusion beyond the baseline capacity of the uncoated scaffold. This finding is consistent with the established role of collagen IV in promoting myoblast differentiation^46,47^. Multinucleation analysis further supported this conclusion, with the collagen IV group showing significantly greater proportions of multinucleated myotubes compared to the laminin/collagen IV and CG groups, and the laminin group also showing greater early fusion in the 2-4 nuclei bin compared to the laminin/collagen IV group (**Fig. 5l**). Myotube orientation analysis revealed modest preferential alignment in the longitudinal plane across all groups, suggesting that myotube organization can be guided by the scaffold anisotropic pore structure (**Fig. S3**).

Critically, the diminished fusion and multinucleation of muscle cells observed in the laminin/collagen IV group compared to the collagen IV group is consistent with the reduced collagen IV functionalization observed in **Fig. 2**, where potential interaction and interference during simultaneous functionalization appeared to limit collagen IV incorporation. Taken together these findings suggest that collagen IV is helpful for myogenic differentiation in this scaffold system, and that its dilution by co-application with laminin compromises its pro-fusion signaling capacity. This has direct practical implications for scaffold functionalization strategies where alternative approaches to simultaneous application of basement membrane proteins may better preserve the individual biological activities of each component and warrants investigation in future work. Laminin functionalization supported myotube formation comparable to the uncoated CG scaffolds across most metrics, consistent with laminin’s established role in maintaining progenitor cell state rather than directly promoting terminal differentiation^43^. The similar performance of the laminin and uncoated CG groups across fusion and multinucleation metrics suggest that while laminin provides important niche signals for progenitor maintenance, collagen IV provides an important pro-differentiation cue in this system.

### CG scaffolds support neural stem cell metabolic activity

As an initial step toward assessing the suitability of our scaffold platform for supporting non-myogenic cell types helpful in skeletal muscle tissue engineering, we cultured primary rat NSCs on aligned CG scaffolds for 8 days. Fluorescence imaging confirmed that NSCs remained attached to the scaffold surface and retained nuclear integrity across all coating conditions throughout the culture period, with positive β-III tubulin staining observed in all groups, potentially indicative of early neuronal differentiation (**Fig. 6a-d**). NSC metabolic activity did not significantly differ between groups for the majority of time points, with the exception of higher activity in the CG group compared to collagen IV and laminin/collagen IV groups on day 4, and higher activity in the laminin group compared to the laminin/collagen IV group on day 8 (**Fig. 6e**). These findings indicate that NSCs remained metabolically active across all ECM-functionalized scaffold conditions, establishing initial feasibility for NSC compatibility with the functionalized scaffold platform, and supporting future investigation into neuromuscular co-culture strategies for VML repair.

**Figure 6.**
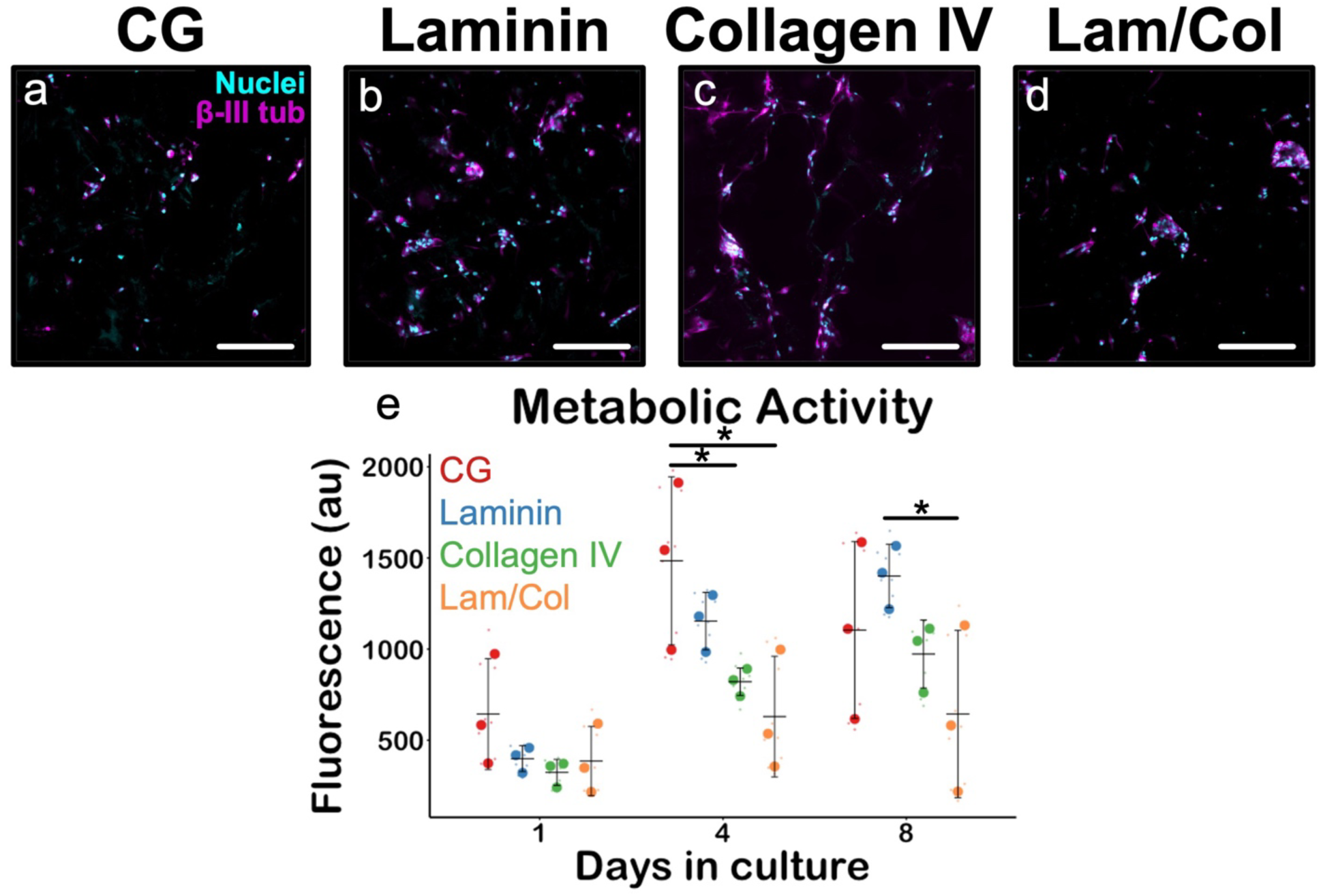
CG scaffolds support NSC metabolically activity. (a-d) Representative fluorescence images of NSCs cultured in scaffold groups following 8 days of culture, stained for β-III tubulin (magenta) and DAPI (blue). (e) Metabolic activity assessed by alamarBlue assay at days 1, 4, and 8 of culture. Scale bars: 200 µm. *N* = 3 scaffolds. *: p < 0.05.

## Conclusions

This work establishes a CG scaffold platform combining an anisotropic pore structure with basement membrane protein functionalization to capture key skeletal muscle structural and compositional cues. We show that aligned CG scaffolds can be successfully functionalized with laminin, collagen IV, and their combination while preserving pore architecture and achieving stable protein retention over 7 days. Collagen IV functionalization produced the greatest increase in hMPC fusion and myotube maturation, consistent with its established role in promoting myogenic differentiation. Myotube orientation analysis further revealed modest topographical alignment along the scaffold longitudinal axis, confirming the ability of the anisotropic pore structure to guide myotube organization. Preliminary neural experiments demonstrated NSC attachment and sustained metabolic activity across all scaffold conditions, establishing initial feasibility for use of this cell type with the CG scaffold platform. Ultimately, this work sets the foundation for future neuromuscular tissue engineering strategies and *in vivo* validation in VML animal models.

## Acknowledgments

The authors thank Professor George Christ for his contribution of the human-derived muscle progenitor cells. We thank Roland Buchanan for his expertise and fabrication of the freeze-drying molds. We thank Professor Kyle Lampe for his knowledge and advice regarding the optimization of neural cultures. We thank Dr. James Gentry for his helpful discussion and work on the myotube metric Python pipeline. We thank Annabelle Hendrickson, Jeremy Ortmann, and Megan Lee for their support, critical evaluation, and review. SEM images were acquired at the University of Virginia Nanoscale Materials Characterization Facility (NMCF). This work was supported by the NIH (R01AR078866, R35GM162024). The content is solely the responsibility of the authors and does not necessarily represent the official views of the National Institutes of Health.

