## Supplemental Information for "Aligned basement membrane-modified collagen scaffolds for skeletal muscle tissue engineering"

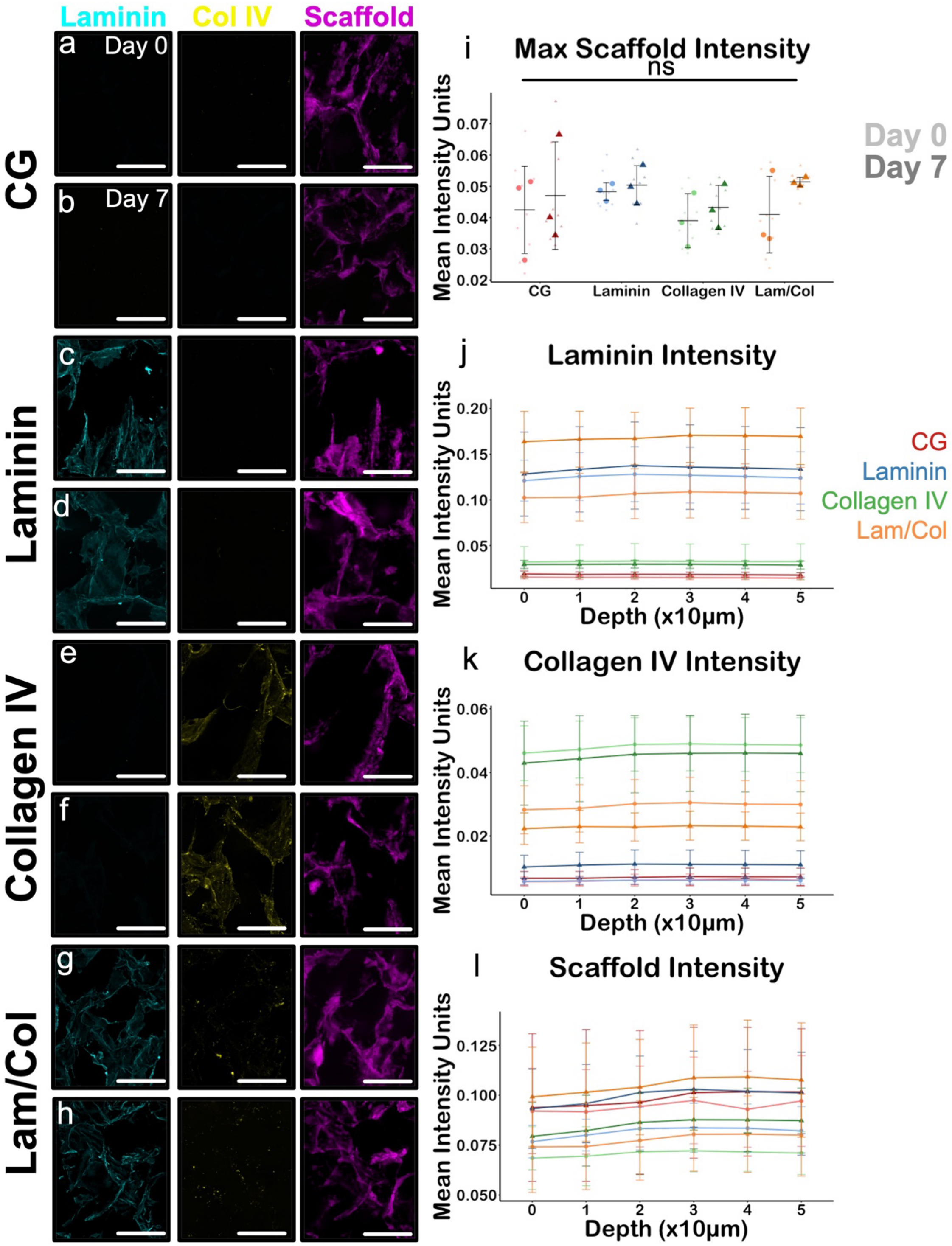


**Figure S1. ECM proteins remain stably associated within the scaffold over 7 days.** (a-h) Fluorescence images of ECM proteins, separated by channel, of each group show positive staining and retention within the scaffold. (i) Quantification of fluorescence intensity demonstrates no significant reduction in scaffold signal across groups between day 0 and day 7. (j) Laminin, (k) collagen IV, and (l) scaffold staining remained constant throughout the scaffold. Scale bars: 200 µm. *N* = 3 scaffolds per group. *: $p < 0.05$.

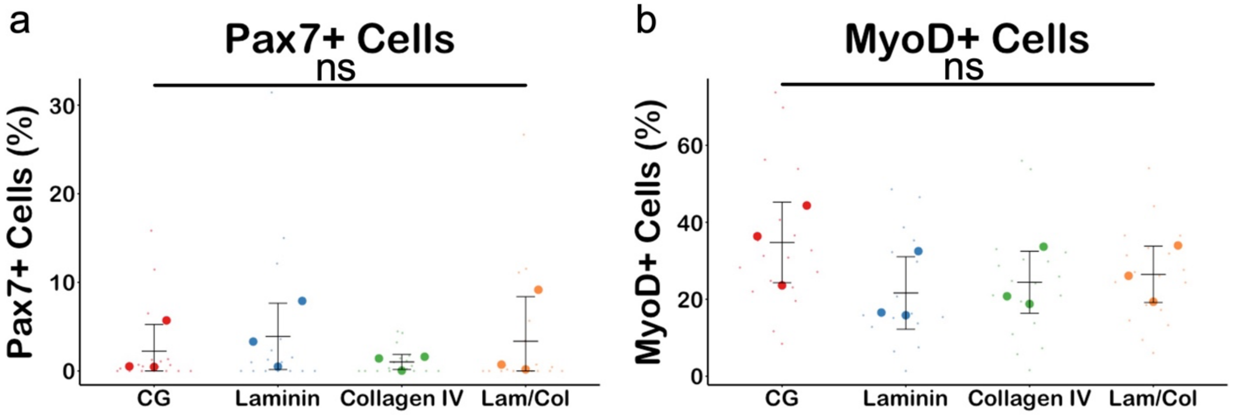


**Figure S2. ECM protein coating does not significantly affect satellite cell maintenance and differentiation.** (a) Relative proportions of Pax7+ cells and (b) MyoD+ cells remain consistent across different ECM functionalization conditions. *N* = 3 scaffolds.

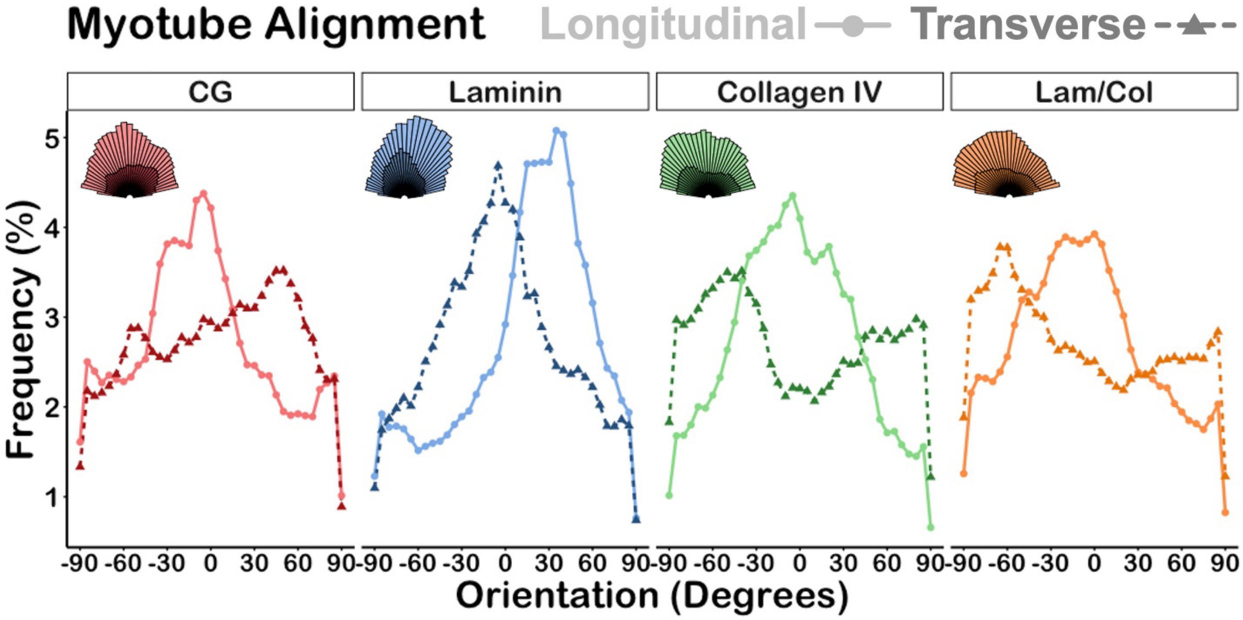


**Figure S3. Myotube orientation shows modest preferential alignment in longitudinal plane.** Quantification of MHC+ myotube orientation reveals modest preferential alignment in the longitudinal plane across all scaffold groups. *N* = 6 scaffolds.
